# Machine learning model determines optimal coast redwood restoration sites in Santa Clara County, California

**DOI:** 10.1101/2025.03.18.644027

**Authors:** Richard B. Lanman, Christopher Potter

## Abstract

Many governmental and non-governmental organizations are planting trees, often in developing countries where costs are lower, to offset carbon emissions in industrial countries and slow global warming. These efforts often fail to achieve carbon sequestration goals, frequently related to the selection of unsuitable planting sites, use of tree species with inadequate biomass or vulnerability to wildfire. Reforestation of the world’s tallest trees, which attain heights above 80 m and standing biomasses orders of magnitude greater than other trees, could optimize removal of atmospheric carbon. Wildfires have consumed almost 30% of California’s forests since 2000. To optimize carbon sequestration by tree planting in Santa Clara County, California, we selected coast redwoods, a tree species that is highly resilient to fire, rapidly accumulates the largest biomass of any tree in the world, has exceptional longevity, and is historically native to the westernmost portion of the county, where they support a unique ecosystem. Because historical redwoods range maps are conflicting, and because global warming may change the range of suitable habitat, we developed a machine learning model to determine optimal habitat available for reforestation. The optimal habitat identified included the current and known historical coast redwood forest and excluded lands with no historical records of redwoods, validating model accuracy. The opportunity to capture and store atmospheric carbon dioxide (CO_2_) is significant, as the model found 33,969 ha (131 mi^2^) of optimal redwood habitat, of which nearly 77% [26,051 ha (101 mi^2^)] currently has no conifer cover. Restoring the historical coast redwood forest in this single county alone could sequester 2.3% of the entire State of California’s 2020 carbon emissions. Similar studies should be conducted in ocean-adjacent counties which likely have order of magnitude greater optimal coast redwood planting areas than inland Santa Clara County, with correspondingly greater potential carbon sequestration impacts.

## INTRODUCTION

Planting trees to sequester atmospheric carbon may potentially provide over one-third of the climate mitigation needed to slow or stabilize global warming (1). In addition, re-established forests cool regions by 1°–2°C compared to neighboring grasslands and croplands (2). Recent innovations that catalyze forest restoration at reduced costs are now in practice (3). As a non-signatory to the Kyoto Protocols, American landowners do not benefit from international emissions trading, but regional forest carbon credits programs were pioneered in California with some projects (but not all) demonstrating additionality, or reducing net atmospheric carbon more than would have occurred naturally, (4–6). As of 2020, approximately 25% of global annual anthropogenic carbon emissions were offset by forest ecosystems (7). However, the latter are continually diminished by timber harvest and development, with deforestation today the third largest cause of global CO_2_ emissions globally, accounting for up to one-fifth of the total (8, 9). California, however, is a special case, where wildfires have reduced forested land area far in excess of timber harvest and conversion of land to agriculture, with 29% of tree cover burned since 2000, equivalent to 2.83 million hectares lost (10, 11). Wildfire frequency in California has doubled since the 1980s, and further increases in megafires, which reach and consume forest canopies, are expected (12, 13). In just the last decade, 20% of California’s forest carbon buffer pool, meant to insure against unforeseen losses to forest carbon offset projects for a century, was destroyed by wildfires (14). Wildfires threaten California’s carbon offset credits program, as industry purchasers of carbon credits are permitted to emit greenhouse gases while the forest stocks of owners who were awarded credit payments fall below their baselines (6, 15). Therefore, we submit that California reforestation and afforestation initiatives in California should focus on fire-resistant tree species.

Two native tree species in California, both in the cypress family (*Cupressaceae*), possess the twin qualities of resilience against fire and exceptionally high biomass combined with being amongst the fastest-growing trees globally, namely coast redwoods (*Sequoia sempervirens*), and giant sequoia (*Sequoiadendron giganteum*) (16–18). Further, these tree species achieve the highest standing carbon stocks of any trees in the world (19, 20), with the coast redwood being the tallest although the giant sequoia having the greatest girth (21, 22). The tallest organism on earth is thought to be a coast redwood standing at a height of over 112 m (23). A single large coast redwood [91 m (299 ft) tall, ∼7 m (23 ft) diameter] was determined to have the same carbon stock as an entire hectare of eastern temperate forest (24). The emergent crowns of old-growth coast redwoods, which tower over surrounding forest canopy, also accumulate complex structures composed of living and dead wood, further contributing to their above-ground biomass, so that their above-ground carbon biomass exceeds that of plantation or second-growth forests of coast redwoods (20,25,26). Both species are adapted to resist California’s frequent fires, possessing unusually thick and resin-free bark on their lower trunks, and both are capable of epicormic sprouting along their trunks post-fire, even after high burn severity canopy fires (19, 27). The combination of resistance to fire and fungal decay also contribute to the longevity of both species, with lifespans exceeding 2,000 years in mature coast redwoods and 3,000 years in giant sequoia (19, 28). Furthermore, resistance to fungal decay contributes to the long-term carbon sequestration of these trees, which may take an additional 1,000 years to decompose after they die (29). However, only coast redwoods are native to our study area, Santa Clara County, California. Surviving coast redwood forests are important biomass sinks for California’s mandated CO_2_ sequestration objectives (30). However, only 5% of the original old-growth coast redwood forests that once covered 890,000 ha (2.1% of California’s total area) remain after decades of logging for lumber (19, 31).

Coast redwood forests grow natively in a narrow band along or near the Pacific Coast from southern Oregon to southern Monterey County, California. They depend on a maritime climate, which provides relatively stable temperature and soil moisture availability (32, 33). Of the variables predicting suitable planting locations for coast redwoods, their high demand for water is most important (32). Within coastal redwood forests of northern California, water use has been estimated at around 600 liters per day for a 45 m tall redwood tree (34). The highest demand for water occurs during the summer months when evapotranspiration from foliage is high (35). Although summer rainfall is rare in the region, the capture of fog moisture by redwood canopies can offset some of the rainwater deficits (32). In a study of redwood tree distribution in three different forest reserves in California, the growth rates and population sizes of coast redwood trees was strongly dependent on proximity to streams and creek beds. Although four factors affecting access to water best predicted redwood tree prevalence: soil water storage, height above nearest stream, fog frequency, and heat load, the influence of these factors varied from one study site to another (33).

In addition to their high biomass and fire resistance, coast redwoods in mature forests provide multiple ecosystem services. Redwoods have been described as a keystone species supporting a unique assemblage of flora and fauna related to their height and amplification of fog-related hydrological inputs to the forest floor (32, 36). They support uncommon or rare plant species in the region such as California huckleberry (*Vaccinium ovatum*) and other epiphytes, including ferns, orchids, lichens, and mosses (37, 38). California spotted owls (*Strix occidentalis occidentalis*) and marbled murrelets (*Brachyramphus marmoratus*), both federally threatened bird species, nest in the canopies of mature redwood forests, including in the Santa Cruz Mountains (39, 40). Redwoods provide shade that helps maintain the cool water temperatures essential for salmonid spawning and rearing (41). Also, the large woody debris from fallen redwoods shelters both young salmon and steelhead trout *(Oncorhynchus spp.*) and creates deep perennial scour pools essential to their dry season survival in the region’s numerous intermittent streams (42). Litter and soil created from the decomposition of redwood organic matter falling from live tree canopies provides high-quality mulch and compost for native plants (43).

The North Santa Clara Resource Conservation District (NSCRCD) is an independent special district of the state of California, whose mission is the conservation of soil, water and wildlife in north Santa Clara County. Because of coast redwoods’ high biomass, longevity, rapid growth rate, and resistance to fire and decay, NSCRCD identified reforestation of this tree species as a priority to promote carbon sequestration and mitigate global warming. Hence, the goals of this study were to (1) identify suitable planting sites for coast redwoods in Santa Clara County, California and (2) develop an optimal redwood habitat model which could be re-applied and leveraged for the other 12 counties in the trees historical range, or even further north. Although Santa Clara County is not adjacent to the Pacific Ocean, coast redwoods are native to its westernmost portion, on the inland slopes of the Santa Cruz Mountains (44, 45). In neighboring coastal Santa Cruz County, Big Basin Redwoods State Park (BBRSP) is renowned for its ancient coast redwood trees, many of which are over 100 m (300 ft) tall and 1,000–2,500 years old (28). Many of the BBRSP redwoods survived a high-severity wildfire, the 2020 CZU Lightning Complex Fires (CZU Fire) in Santa Cruz County, even though their canopies were burned (46, 47).

As a precursor to the present study, in 2017 the Turtle Island Restoration Network’s 10,000 Redwoods Project undertook a novel effort to identify habitat suitable for coast redwood restoration in Marin County, California (48). For their model, they used geographic information service (GIS) spatial analysis modeling that utilized seven key factors conducive to redwood growth based on an extensive literature review. We built on this prior work via application of sophisticated machine learning (ML) algorithms to determine key factors and weights predictive of optimal coast redwood planting areas. Specifically, Random Forest (49) ML was used for development of optimal habitat mapping based on key climate, hydrologic, and topographic predictors; this ML approach has the ability to be trained with existing nearby mature coast redwood cover areas in order to account for local rather than range-wide factors. Although redwoods had a much broader range prehistorically, their current narrow distribution along the California coastal belt suggests that local factors are important to redwood growth. These include generally stable microclimate regimes without extremes of heat or frost, low wind exposure, lack of sea salt aerosol, either fog or high soil moisture presence, and alluvial soils typical of valley bottoms (50). In a study comparing three different mature groves spanning the middle latitudinal third of coast redwoods range, the factors influencing redwoods presence varied significantly. At BBRSP the best predictor of redwoods presence was soil water storage, followed by height above stream, and then fog; however, on Mt. Tamalpais further north, fog was the best predictor, followed by soil water storage, temperature and height above stream last (33). These findings highlight the importance of local factors in predicting optimal redwood habitat. Machine learning models enable utilization of local input variables and also enable both regression prediction and map-based classification (51). Finally, machine learning models ML can be continuously updated based on new data and learning, such as climate change, improving the effectiveness of initial models over time (52, 53). However, ML models have not yet been applied to determination of optimal habitat for planning coast redwood reforestation.

## METHODS

### Study Area

The study area was the Santa Clara County, California portion of the Santa Cruz Mountains, whose midpoint is a summit named The Peak, elevation 913 m at coordinates 37.219 N, 122.072 W. The Santa Cruz Mountains consists of two northwest to southeast trending ranges bisected by Los Gatos Creek, which was dammed to form Lexington Reservoir. The Spanish named the northern range the “Sierra Morena” or dark mountains, and the southern range the “Sierra Azul” or blue mountains (54). Santa Clara County is relatively unique as 31% (104,248 ha) of its total area (337,736 ha) are protected open space lands (55, 56). Its juxtaposition of a major metropolitan area with extensive conserved lands is attributable to over 120 years of land preservation initiated by several governmental and non-governmental organizations, beginning with the Sempervirens Fund in the Santa Cruz Mountains in 1901 (57).

### Historical Distribution of Coast Redwoods

Western Santa Clara County includes primarily the inland slope of the Santa Cruz Mountains to the summit divide although it also includes minor components of the range’s Pacific slope. The Santa Cruz Mountains lie approximately three-quarters of the way down the 750 km Pacific Coast distribution of coast redwoods, which stretch from Curry County in southwestern Oregon to southern Monterey County in California. The range was first described as historical redwood habitat in the 1776 De Anza Expedition by Father Pedro Font: “*… all the way from the vicinity of the Arroyo de las Llagas* [Gilroy, California] *there runs clear to the Punta de Almejas* [Pacifica, California] *a very high range, most of it thickly grewn with cedars* [redwoods]*…”* (58). As inferred from the locations of past redwood sawmills, the redwood forest historically extended down the inland side of the Santa Cruz Mountains at least to the foothills at the western edge of the Santa Clara Valley floor (45) (Fig 1). These lumber mills were typically based in the forest, given the difficulty hauling colossal redwood logs.

**Fig 1.**
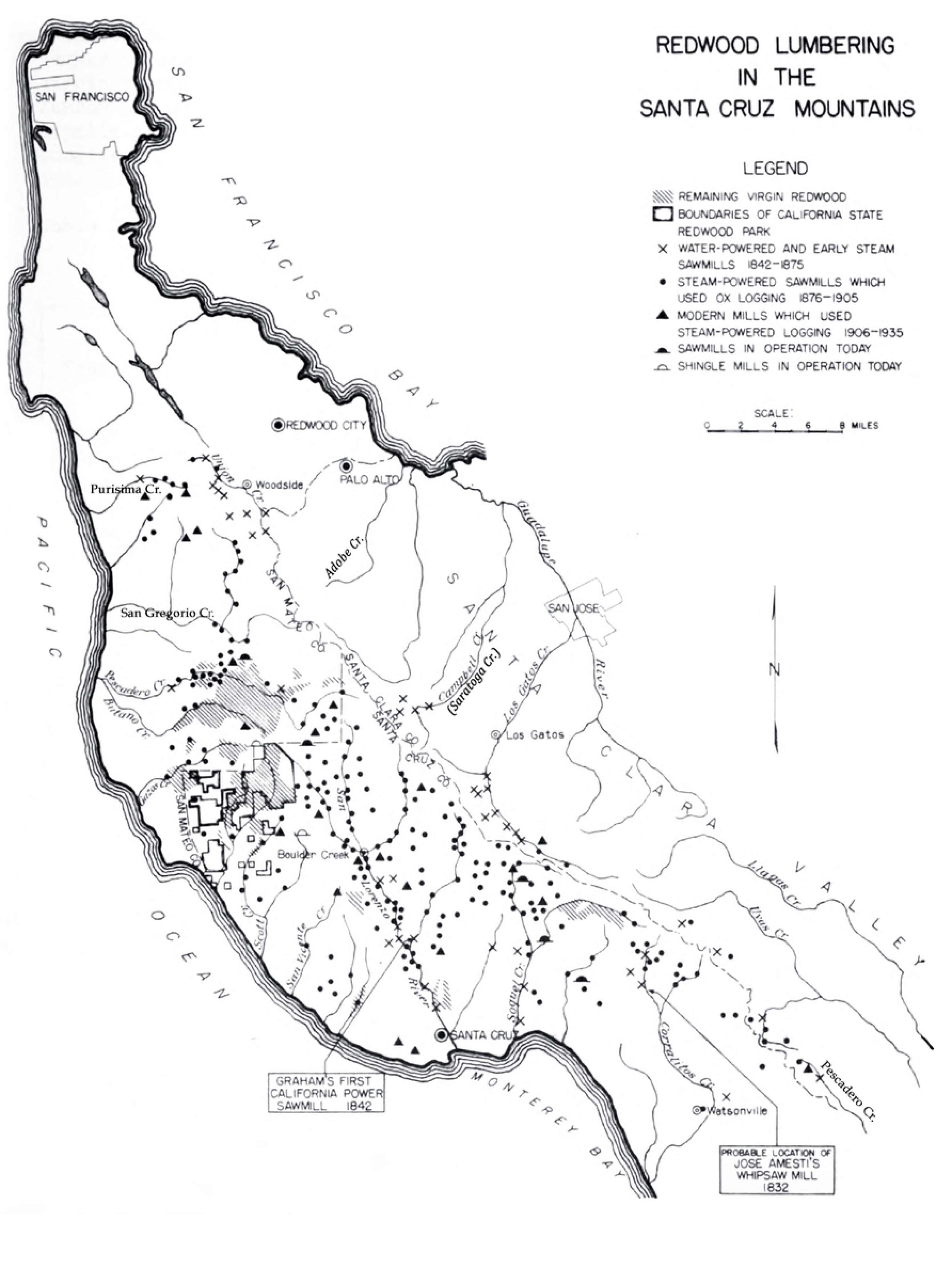
U.S. Forest Service map of historical redwood lumber mills reflects past extent of redwood forest extending across the Santa Cruz Mountains from coastal San Mateo and Santa Cruz Counties, and to inland Santa Clara County (which includes the western edge of the Santa Clara Valley). Callouts highlight the earliest known water-powered sawmills in the region in 1842. Map adapted from Jensen 1939.

NSCRCD initially sought to use maps of the historical range of coast redwoods as a guide to restoration, however, three issues challenged this approach. First, precise records of redwoods’ historical distribution in the region were incompletely established because logging began long ago in the 1770’s. A leap in demand for lumber in the San Francisco Bay Area related to the California Gold Rush resulting in cessation of logging records on the inland side of the Santa Cruz Mountains by the 1850’s, suggesting that the entire old-growth redwood forest in Santa Clara County had been removed (59). Logging continued on the coastal side of the range until deforestation was nearly complete in most of the Santa Cruz Mountains region by the 1870’s (60) (Fig 2). Today, second-growth redwoods, often arranged in fairy rings (the clonal sprouts which grow from the circular periphery of felled parent trees) can be found at elevations as low as 100 m (328 ft) along Pescadero Creek in the southern Santa Cruz Mountains in Santa Clara County or 240 m (787 ft) at the intersection of Redwood Gulch and Stevens Canyon Road on the inland side of the northern Santa Cruz Mountains. These findings are consistent with Jensen’s 1939 map of historical sawmills, indicating that historical redwood forest extended inland from the Santa Cruz Mountains nearly to the floor of the Santa Clara Valley (45). Second, historical distribution maps may not accurately depict suitable coast redwood habitat today, as the state has warmed 1.4° C–1.5° C since the late nineteenth century and is currently warming at an accelerated rate (35, 61). Thirdly, maps of the historical depictions of redwood forests in the region are somewhat conflicting, particularly on the extent of the forest further inland onto the Santa Clara Valley floor (62)(Sonia Morris and Joanna Nelson, Save the Redwoods League, personal communication; Robert Van Pelt, University of Washington, personal communication).

**Fig 2.**
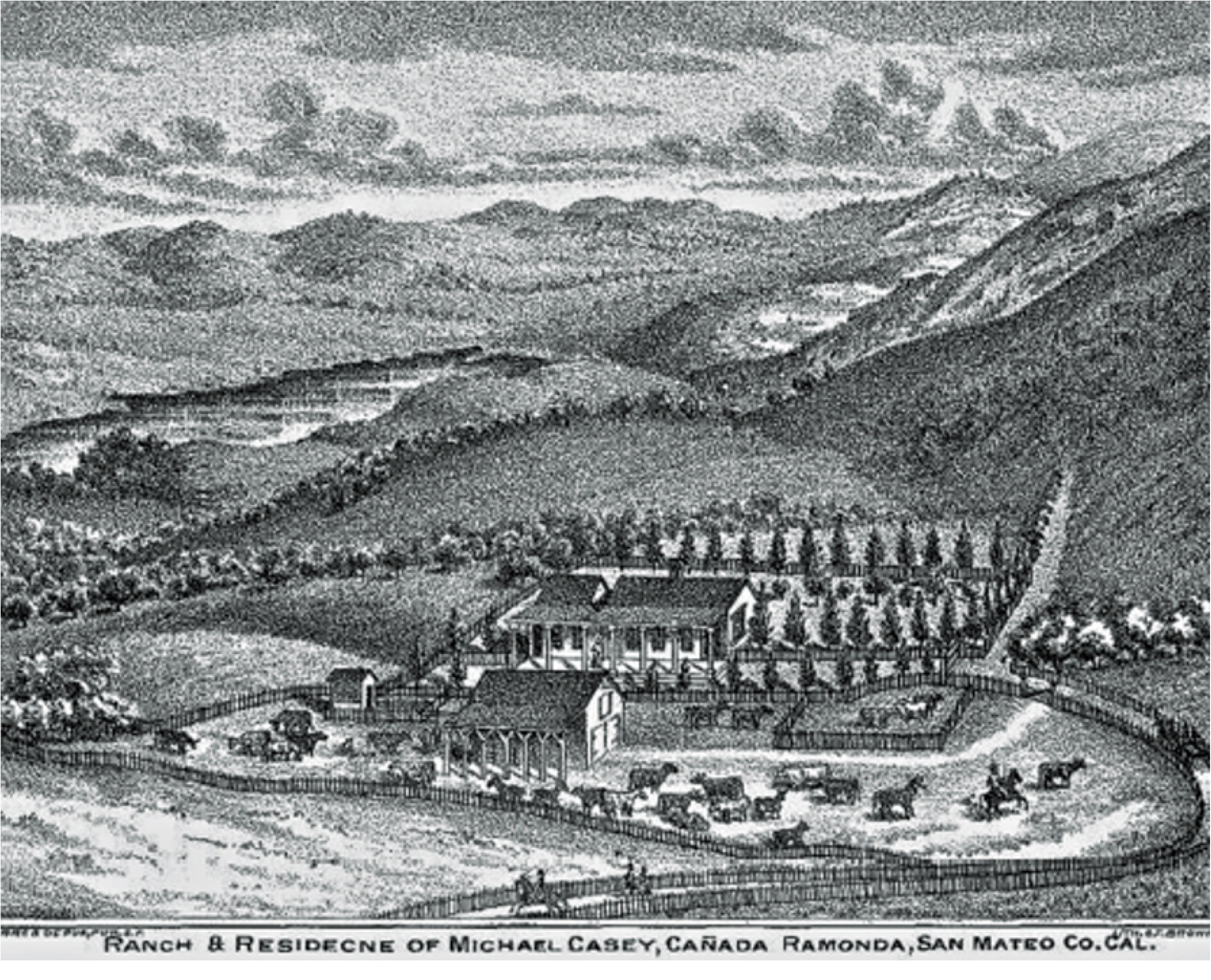
Lithograph survey of 1878 highlights the complete deforestation from timber harvest of the inland side of the Santa Cruz Mountains by the 1870’s (63). Overview from the historical Casey Ranch near today’s California State Route 92 looks east over the Cañada Raymundo, the valley including Upper Crystal Springs Reservoir. Today, second-growth coast redwood forest thrives today on the inland side of the Santa Cruz Mountains from the ridgetops down nearly to the reservoir.

### Experimental Plots and Survival of Coast Redwood Seedlings

We evaluated survival of experimental plots of one-year-old redwood seedlings after 19 months of staging/further aging in marginal habitat and 12 and 24 months after transplantation to optimal redwood habitat. Two hundred fifty one-year-old coast redwood seedling plugs were obtained from the California Department of Forestry and Fire Protection’s (CAL FIRE) L. A. Moran Reforestation Center (64). Trees were selected from Zone 079 of the California Seed Bank, a long-term repository of native conifer seeds sorted and stored by elevation and local climactic/physiographic zone to match historical redwood habitat at elevation 396 m (1,300 ft) in western Santa Clara County. These were initially planted in two equally divided plots of 125 trees each in May 2021 in marginal redwood habitat at elevation 38.4 m (126 feet). One plot was in full sun and one in afternoon shade. Both plots were watered monthly from May to October 2021 but received no supplemental water from months eight to 19. Tree survival was evaluated at months seven and 19. At the end of this period, November 2022, the 2.5-year-old trees were transplanted approximately 10 m apart along a 500 m reach of an ephemeral stream, 5–25 m from thalweg, on 5° to 25° slopes in partial shade in oak (*Quercus* spp.) woodland, and at elevation 400–500 m (1,312 ft to 1,640 ft). The 10 m spacing was based on mature coast redwood forest spacing of 25 to 90 trees per hectare (65), which if 90 trees per acre were evenly spaced, would place the trees 10.6 m (34.78 ft) apart. We continued to evaluate cumulative tree survival at one and two years (November 2023 and 2024) post-transplantation.

### Machine Learning Model

Polygon boundary shapefiles for BBRSP, Henry Cowell Redwoods State Park, and Forest of Nisene Marks State Park (SP) in Santa Cruz County California were used as coast redwood habitat training areas for ML prediction and classification in this study (Fig 3). Within each of these three SP areas, 10,000 sample locations were randomly selected in ArcGIS as training/testing points to input into ML prediction algorithms of SciKit-learn library written in the Python programming language (66). Scikit-learn features various classification and regression algorithms including decision trees, support vector machines, Random Forest, and K-means nearest neighbor, all operating with the Python libraries NumPy and SciPy. Among the available Scikit-learn ML methods, we selected Random Forest (49) for model development of optimal coast redwood forest habitat, because it has the ability to perform both robust regression prediction and map-based classification (51). Random Forest is generally considered to be an improved extension on classification and regression trees (CART) (67). Random Forest models operate as follows: first, the algorithms computationally “grow” a forest of ntree trees. For each tree from 1 to ntree, a sample of size N is taken from the dataset with replacement (bootstrap) to grow the tree. A selection of m variables, independently for each node tree, is made, and the tree is split at each node by determining which variable will create the highest proportion of homogenous classification using Gini impurity criteria. In a random forest model, Gini impurity measures how likely a randomly selected data point is to be misclassified (49). Trees are grown until the nodes can no longer be split, unless otherwise specified with a max_depth variable to prevent overfitting of the data. For model training, 90% of the data points were selected while the remaining 10% of data points were reserved to create the “testing” data set, used to unbiasedly validate the model’s fit and accuracy based on the training model result.

**Fig 3.**
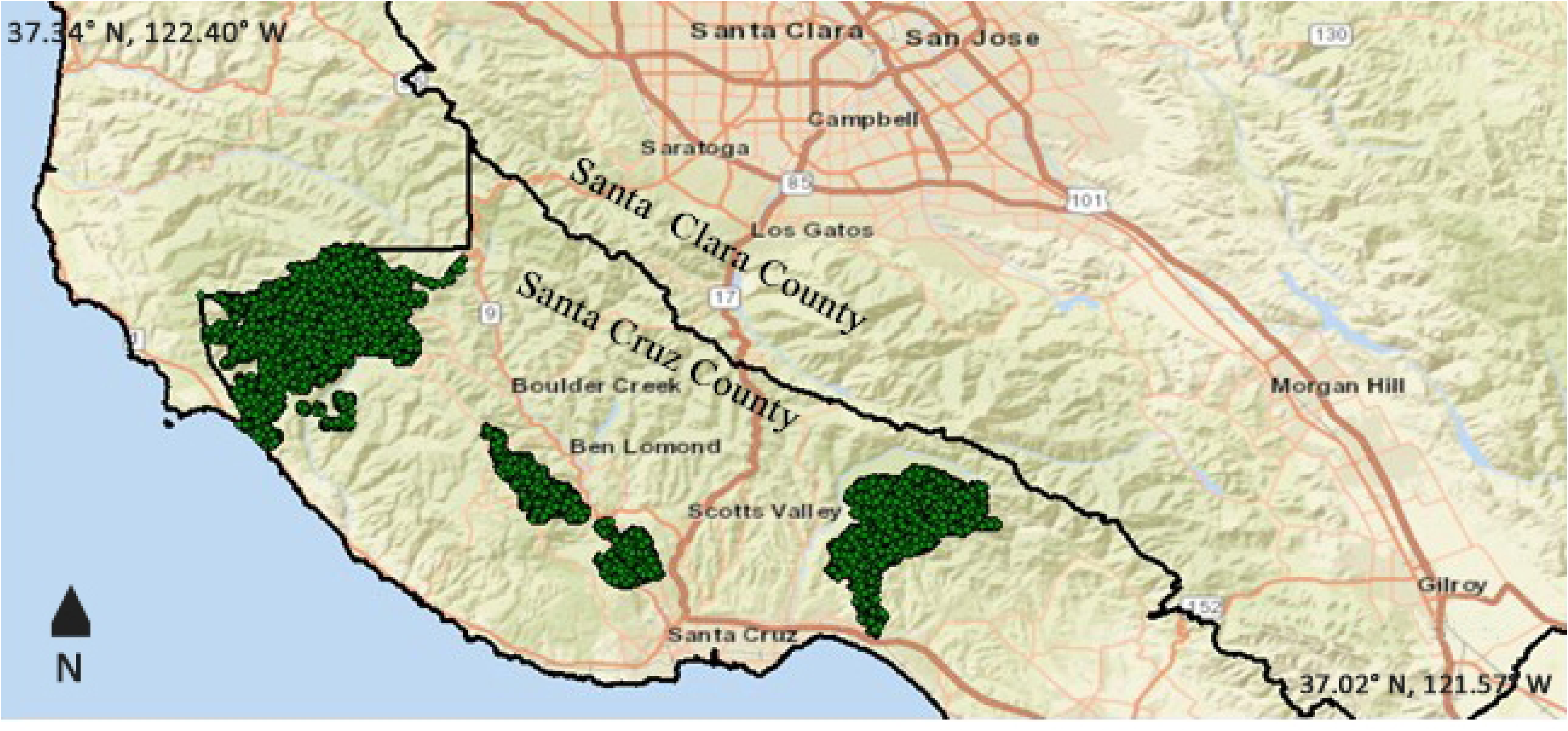
Map of three nearby redwood state park areas with 10,000 sample locations each, randomly selected as training/testing points to input into machine learning (ML) prediction algorithms. GPS coordinates reflect northwest and southeast corners of the map, respectively.

The California streams line shapefile within each of the three redwood SP was used to determine the distance (m) of any location to the nearest stream course (DTNS), which, in an inland county like Santa Clara County with minimal coastal fog, would be essential to provide adequate soil moisture. Therefore, DTNS was set as the predicted diagnostic variable to determine optimal redwood habitat ratings for all locations in SCC using SciKit-Learn ML classification methods. The largest coast redwoods prefer broad, alluvial valleys with lower slopes (16)so we assumed that optimal redwood habitat should be on slopes of 30° or less. Both DTNS and slope are key and independent factors predicting access to soil water stores and are related to height above the nearest drainage (HAND), which is correlated to the depth of the water table (68), by the equation DTNS = HAND/tan(θ) where θ is the average slope angle. Interpolated height above stream (IHAS), a measure equivalent to HAND, was a key predictor of redwood presence in BBRSP (33). In that study the maximum IHAS (HAND) for coast redwood habitat for BBRSP was 150 m above stream bottoms (Francis et al. 2020). By limiting DTNS to a maximum of 150 m in our study, we selected a more conservative measure of soil moisture because for DTNS of 150 m and slope angles of 30° or less, HAND would be 87 m or less. Therefore, we set a distance of 150 m in the DTNS layer to be equivalent in the Random Forest prediction model to the highest ranked value result for “optimal” redwood planting habitat locations.

The following independent variable values for all training and testing points were extracted from six 30-m (Landsat image resolution) raster layers and input to the SciKit-Learn ML prediction script: elevation, slope, aspect, land surface temperature (LST) in July 2020, plant water use (ET flux in mm per year in 2019), and United States Department of Agriculture (USDA) soil type properties for depth to bedrock, moisture holding capacity, and parent material.

Elevation, slope, and aspect layers were extracted from the United States Geological Survey 1 arc-second (30 m) resolution data layers (USGS 2023). LST in July 2020 (pre-CZU Fire) was estimated from Landsat satellite brightness temperature image maps regridded to 30 m resolution (Potter and Alexander 2021). Plant water use in 2019 (pre-CZU Fire) was derived from the OpenET platform (openetdata.org), which uses inputs from Landsat, Sentinel-2, GOES, and other satellites, along with weather station data and various weather models (ReVelle et al. 2021). Soil types were mapped and gridded to 30-m resolution from the USDA Soil Survey Geographic Database (Soil Survey Staff 2023).

## RESULTS

Random Forest results for prediction of DTNS-based redwood habitat ratings within the three redwoods SPs in Santa Cruz County returned a testing *R^2^* value of 0.59, and a root mean squared error of 142 m for the predicted DTNS values. This ML model was designed to rely on six spatial variables (Table 1) to account for the influence of topography, micro-climate, water use, and soils on redwood forest habitat. These variables represent a comprehensive set of predictors and together resulted in a high model accuracy in the “testing” subset specified.

**Table 1.** Importance values of each of the input variables for prediction of the distance to nearest stream (DTNS) in redwood SP training areas shown in Fig 3.

| Input Variable | Importance Value |
| --- | --- |
| Elevation | 0.28 |
| Aspect | 0.18 |
| Land Surface Temperature | 0.16 |
| Water Use | 0.14 |
| Slope | 0.12 |
| Soil type | 0.12 |

The Importance Values of each of the input variables for this Random Forest (Table 1) showed that Elevation had the highest value, followed by Aspect and then LST and Water Use. Importance Values indicate the predictive weight of each input variable and are computed as the mean and standard deviation of accumulation of the impurity decrease within each tree. Mean decrease in accuracy is considered the decrease in model accuracy from permuting the values in each feature (49). It is worth noting that, in theory, the range of Importance Values is 0 to 1. However, in practice the range will be considerably lower, because Random Forest randomly picks subsets of the testing data, so there is a good chance that all features were used in a split input format.

These prediction accuracy metrics from ML were judged to be high enough to proceed with the redwood habitat classification model for Santa Clara County using the same six input variables, all gridded at 30-m resolution. To set a threshold for “optimal” redwood planting habitat, we limited DTNS for slopes 30° or below to 150 m to ensure that height above nearest drainage of 87 m was never exceeded and set as equivalent to the Random Forest ranked class value result of 25 or lower in our ML classification analysis and this class was therefore mapped as optimal habitat (Fig 4).

**Fig 4.**
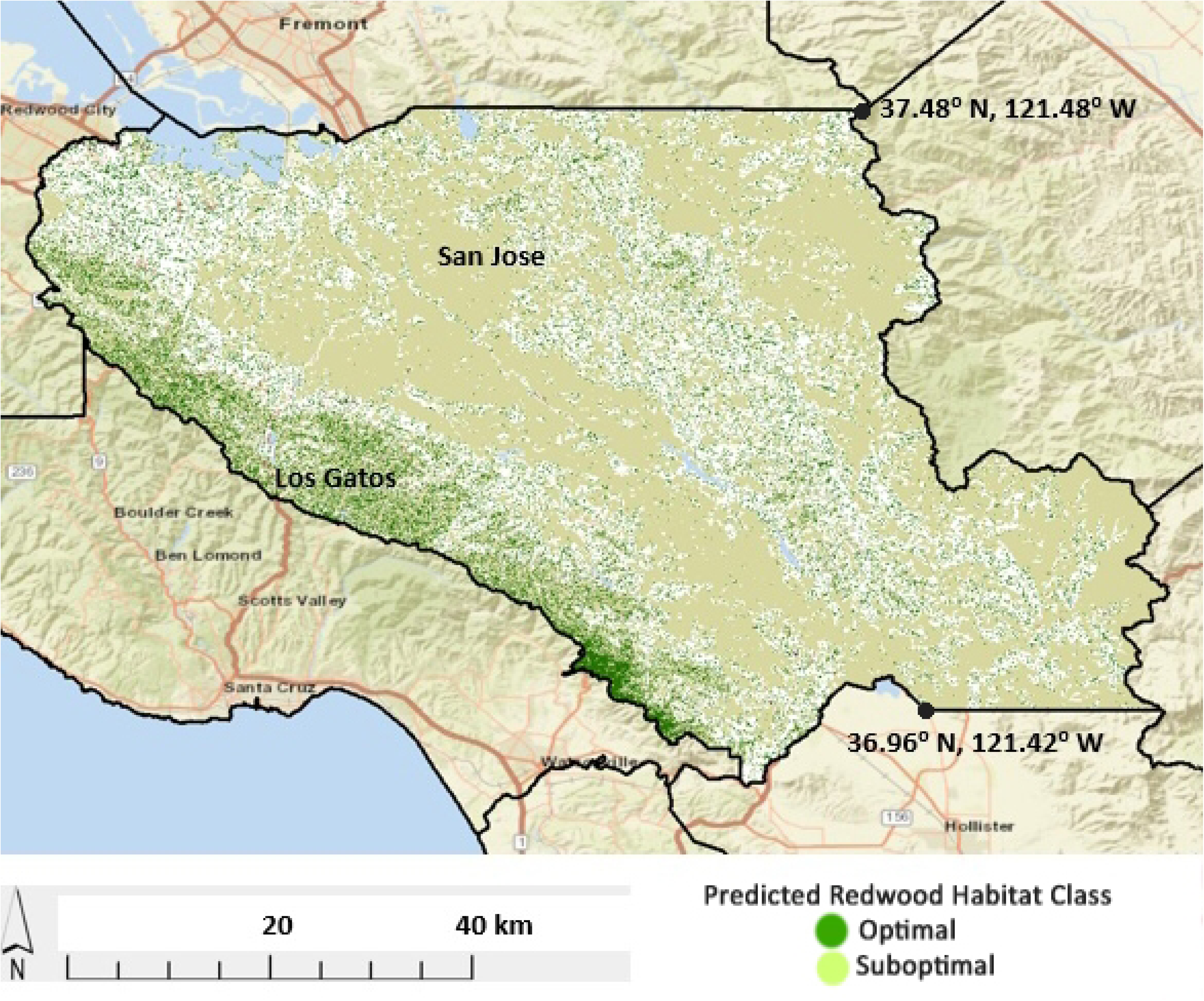
Map output from the redwood habitat classification model for Santa Clara County. Dark green pixels are the “optimal” redwood planting locations and light green pixels show the “suboptimal” locations. As expected from historical maps, the optimal habitat clusters in the western portion of Santa Clara County on the inland side of the Santa Cruz Mountains.

These optimal redwood habitat areas were located in the westernmost portion of Santa Clara County and covered 33,969 ha (83,940 acres), comprising 10% of total County area. “Suboptimal” redwood planting habitat locations in Santa Clara County were mapped in the DTNS-based ranked class value of 26-50, whereas all other ranked class values were considered marginal as redwood planting habitat. The suboptimal redwood habitat areas were more thinly distributed and found across locations across the County at higher elevations and/or close to riparian corridors. Suboptimal redwood habitat covered 420,343 acres 170,107 ha (83,940 acres), 51% of Santa Clara County area.

The densest coverage of optimal redwood habitat was predicted around Mount Madonna County Park near the southern borders of Santa Cruz and Santa Clara Counties. Other dense coverage areas for optimal redwood habitat were identified further north and west, such as in and around Sierra Azul Open Space Preserve and Sanborn County Park. Further to the east, moderately dense coverage areas for suboptimal redwood habitat were identified along creek-sides near Mount Hamilton Road, 4 km north of Smith Creek and Joseph D. Grant County Park, and in low elevation areas between Coyote Lake and Henry W. Coe County Parks. On a county-wide scale, the optimal elevation zones for redwood habitat in Santa Clara had a median value of 400 m (1310 ft), whereas the suboptimal elevation zones for redwood habitat had slightly lower median elevation value at 344 m (1128 ft). The optimal hillslope aspect setting for redwood habitat in Santa Clara County ranged from 160 to 170 degrees SSE.

Turning to climate-related factors, optimal zones for redwood habitat in Santa Clara County showed a median summer LST value of 33.4° C and a maximum summer LST around 38.5° C, whereas suboptimal zones for redwood habitat showed a median summer LST value of 38.4° C and a maximum summer LST of 50.2° C. Annual water demand in optimal zones for redwood habitat in Santa Clara County showed a median value of 924 mm and ranged down to 890 mm, while water demand in suboptimal zones showed a median value of 540 mm. This implied that local (stream, creek, or irrigation) sources must supply no less than 890 mm (35 inches) of water annually to sustain optimal redwood tree growth in Santa Clara County.

Although USDA soil types and properties were not associated with high Importance Value from the Random Forest prediction of coast redwood cover, there was a notable quantity of optimal habitats in Santa Clara County to favor fine-loamy, mesic Mollisols in the sub-group Ultic Argixerolls (69). These soils have an available water content of between 0.12 and 0.17 to a depth of 1.2 m, favorable to the udic soil moisture regime required by redwoods (16, 50). Other attributes of these favorable soils for redwood tree growth were found to be a pH of between 5 and 6, bulk density of between 1.3 and 1.5, fine sand content of 11%, total sand content of 35%, and clay content of 31%.

We next evaluated the percentage of optimal coast redwood habitat found on already conserved lands (Fig 5) utilizing the ML model input and the California Protected Areas Database (70) (Fig 5). A high 78% of the 33,969 ha (131 mi^2^) of optimal habitat southwest of US Highway 101 and California Highway 85 and northeast of the Santa Clara County border (heavy black line) falls on 26,258 ha (98 mi^2^) of already conserved lands.

**Fig 5.**
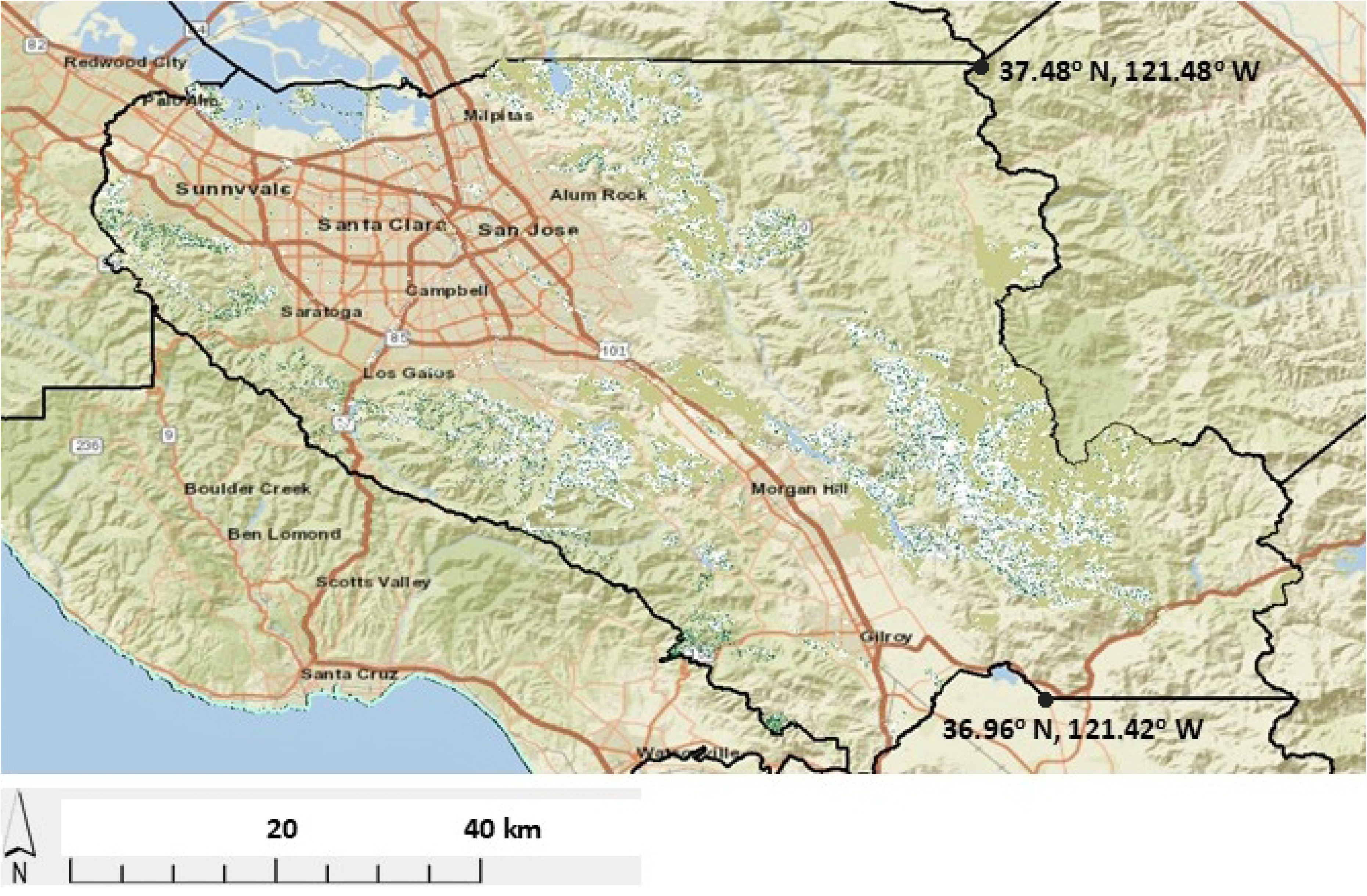
Machine learning map output of optimal and suboptimal coast redwood habitat falling only on conserved lands is shown as dark green pixels on white background. Seventy-eight percent of the optimal habitat southwest of US Highway 101 and California Highway 85 and northeast of the Santa Clara County border (heavy black line) falls on 26,258 ha (98 mi^2^) of already conserved lands.

Next the ML model generated a granular map of optimal coast redwood habitat in the extreme northwestern corner of Santa Clara County, abutting San Mateo County, to demonstrate the model’s ability to identify planting sites at fine (30 m) scale (Fig 6). Higher elevation and proximity to riparian corridors key closely to optimal habitat.

**Fig 6.**
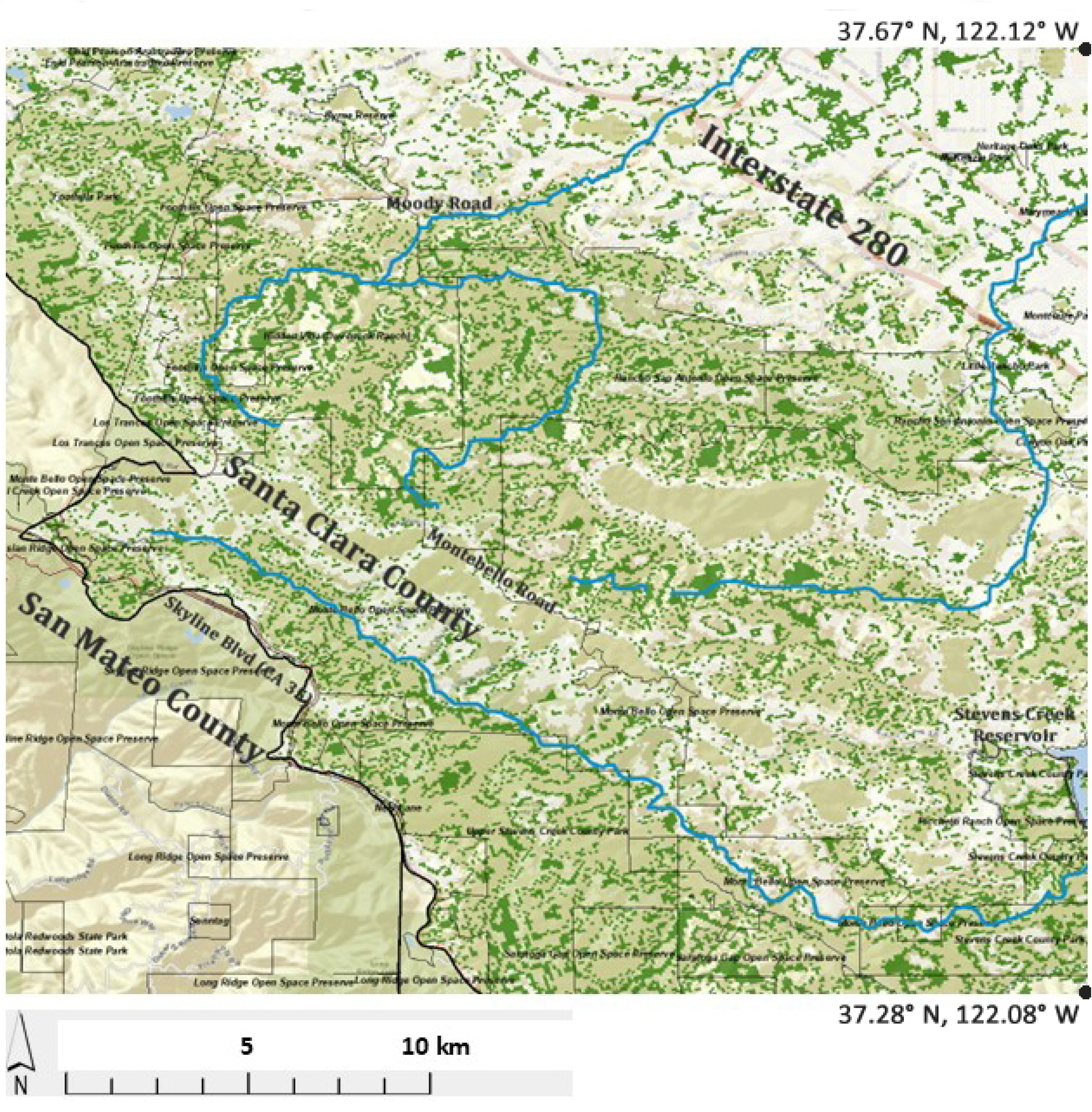
Granular map of optimal coast redwood habit (dark green pixels) illustrates how the ML model may be applied for precise planting site selection. Model results are shown for the northwestern portion of Santa Clara County (and not for San Mateo County which is adjacent on left). Several roads are labeled for orientation as is Stevens Creek Reservoir on lower right. optimal habitat diminishes with lower elevation which falls from southwest to northeast as it approaches Interstate 280 (top right). Optimal habitat clusters with the riparian corridors of three creeks (marked in dark blue): Adobe Creek (top), Permanente Creek (middle right), and Stevens Creek (bottom).

After staging one-year redwood seedlings for 19 months in marginal habitat at elevation 38.4 m (126 feet), survival was 88% in full sun and 95% in afternoon shade. Of note supplemental monthly watering was provided the first six months, but none for the next year. After transplantation of the then 2.5-year-old trees to optimal redwood habitat at 400–500 m elevation cumulative survival was 78% at one year and 74% at two years (Table 2).

**Table 2.**
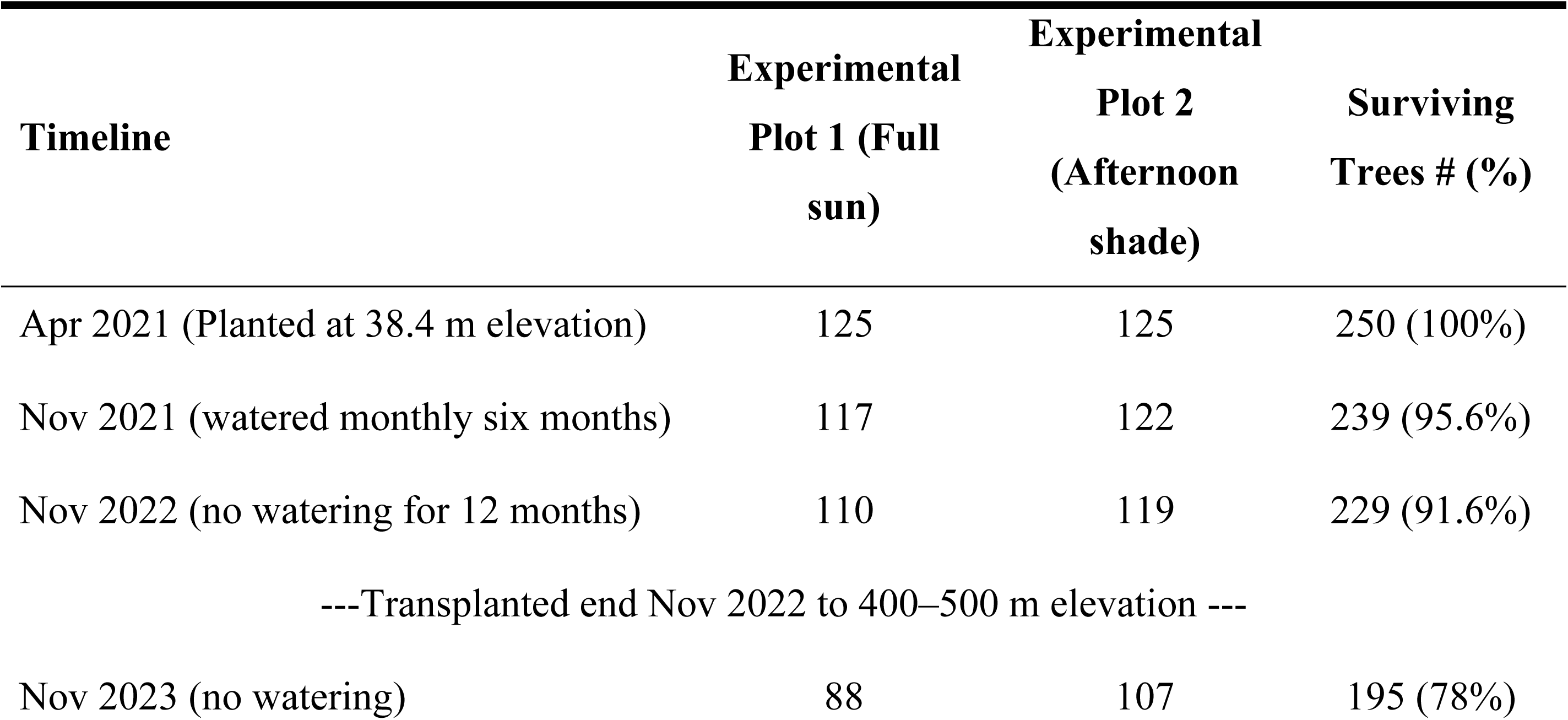
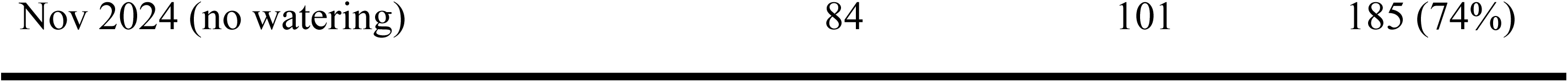
Coast redwood cumulative survival for one-year seedling experimental plots staged locally for 1.5 years in marginal habitat in full-sun or afternoon shade, and then after transplantation of the 2.5-year-old seedlings to model-selected optimal redwood habitat at 549 m elevation for two additional years:

## DISCUSSION

To the best of our knowledge, this is the first application of machine learning to predict suitable planting sites for reforestation and afforestation of coast redwoods. Recently ML applications were broadly recommended for atmospheric carbon sequestration in precision agriculture and forest management, the latter particularly because many tree plantations are started on unsuitable habitat (71). A recent study used ML to evaluate the suitability of planting sites for commercial tree plantations and found that only 0.9% were in highly suitable habitats, 33.2% in medium suitable sites, and fully 66% in low suitability to largely unsuitable sites (53). Our study expands the application of ML to potentially slow global warming by finding optimal planting locations for native coast redwood planting.

The ML model predicted optimal redwood habitat in the Santa Cruz Mountains portion of Santa Clara County where historical sawmill maps indicated the trees once stood and where the few remaining second or third-growth redwood stands exist today, validating the accuracy of the model and its input values for today’s climactic conditions. As important, the ML model did not predict optimal redwoods habitat in areas with no historical records of sawmills. Specifically, it did not predict optimal habitat in the chaparral or oak woodlands in the Diablo Range in eastern Santa Clara County, even where elevations appear suitable, likely because land surface temperatures and water use variables are higher further inland and east. Further, the highest concentration of predicted optimal habitat coincides with where the largest surviving historical redwood groves lie - on those portions of Santa Clara County that extend over the summit ridge onto the Pacific slope, where winter precipitation and summer fog are heaviest. Coast redwoods obtain 19%-45% of their annual water intake from fog, and more importantly, summer growth requires two-to-four-fold higher water intake, making fog critically important in this nearly rain-free season (32). Elevation was the most important input variable in our ML model, reflected by its higher Importance Value of 0.28. The lower temperatures at higher elevation are associated with longer duration of fog, which enables increased precipitation as fog drip in drier summer months (32, 72). There is also reduced evaporative demand from redwood foliage at the reduced temperatures of higher elevations. Water vapor pressure deficits, which increase exponentially with temperature, reduce redwood photosynthesis and growth, and are a major driver of plant mortality in general (35, 73). The two-year 74% cumulative survival rate of transplanted 2.5-year-old redwood seedlings into modeled optimal habitat without further supplemental watering further supports accuracy of the model, and underscores redwoods seedling high shade tolerance and ability to grow in oak woodlands (16). Although the predicted optimal habitat is discontinuous or patchy, this is typical of redwood forest in the southern sub-region south of San Francisco where redwoods require the higher soil moisture found in lower-slope valley bottoms. In fact, redwood forest does not become continuous until north of the Sonoma-Mendocino Counties border (16, 50).

Out of all optimal redwood habitat areas in Santa Clara County mapped by ML models, covering a total of 33,969 ha (131 square miles) county-wide, 26,051 ha (101 square miles) or 76.7% are currently not vegetated by conifer tree cover (74). This is consistent with the complete deforestation of the inland slope of the Santa Cruz Mountains by the 1850’s, coupled with the common practice of redwood stump removal to produce shingles (44, 59). The predominant mode of redwood forest regeneration after lumbering is asexual via clonal sprouts from stumps and root burls; but even these likely succumbed to generations of cattle grazing, vineyards and orchards as stored carbohydrates necessary for repeated sprouting were eventually exhausted.

Lastly, although large quantities of seeds are generated by surviving trees, redwood forest regeneration by sexual reproduction is extremely slow because seed viability is low and of short duration, and seedlings are extremely susceptible to damping off (75, 76). Thus, natural regeneration of the redwood forest in western Santa Clara County, could take millennia (76), leaving the county’s extensive protected ranchlands to succeed from grasslands and chaparral primarily to oak woodlands. “Suboptimal” redwood planting locations that are currently not vegetated by conifer tree cover totaled 161,619 ha (399,369 acres) in Santa Clara County.

Together, the area of potential redwood planting where there is currently little other conifer tree cover occupies 56% of the county-wide land area. However, for the purposes of estimating the potential carbon sequestration by coast redwood planting, and to buffer against continued near-future climactic warming, we have conservatively included only those lands determined as optimal redwood habitat.

Carbon sequestration accrues rapidly after planting redwoods as young trees grow more quickly than almost any other native California conifers (17). Although Monterey pine (*Pinus radiata*) generally grows faster, its short lifespan of 80–90 years makes it relatively unsuitable for long-term carbon sequestration. Young redwood seedlings attain their high growth rates of 0.6 to 2.0 m (2.0 to 6.5 ft) per year at ages of four to 10 years (16). At ages 21–25 years, a redwood plantation in Scotia, Humboldt County, California accumulated wood volume faster than any other tree species, especially noteworthy as no improved seed sources were used and no other management except thinning was employed (77). As the optimal habitat currently occupied by grasslands, chaparral and oak woodlands is replaced by redwoods forest, the carbon in the pre-existing plants and trees would be released. However, a single large coast redwood has the same carbon stock as an entire hectare of average-sized trees, so any loss of pre-existing plant carbon stocks would be rapidly replaced (24). Although young redwoods are more shade tolerant than most conifers, full sun is necessary for rapid growth and may not occur until the above forest canopy is breached. Once emergent, rapid growth continues throughout the redwood’s long life. Dominant young-growth trees on good sites are 30.5 to 45.7 m (100 to 150 ft) tall at 50 years, and 50.3 to 67.1 m (165 to 220 ft) at 100 years (16). The range of sequestration capacity for atmospheric CO_2_ greenhouse gas (GHG) captured in aboveground forest biomass of old-growth (primary) and planted (secondary) redwood stands in historical range in California was estimated at between 400 and 900 Mg ha^-1^ (35). Secondary growth redwood data sets were compiled for this range of biomass storage potential, covering an average of 15 years and a range of 5–34 years of growth. Converting the acreage of optimal redwood planting area in Santa Clara County using these tree biomass data (35), the total storage potential of carbon (roughly 50% by weight of plant biomass) is estimated at between 5.21 and 11.72 (x 10^6^ Mg) million metric tons (MMT) (5,732 MT acre^-1^) of carbon sequestered from the atmosphere in aboveground tree biomass over the next 20–30 years. To put these estimates of forest carbon sequestration into a broader context, this could offset an average of 2.3% of total annual GHG emissions for the state of California in 2020, which were approximately 369 MMT CO2 equivalents (78).

The modeled potential to sequester carbon by planting redwoods in optimal habitat is intentionally conservative by limitation of DTNS to within 150 m on slopes ≤ 30° to ensure height above nearest drainage ≤ 87 m; and further by exclusion of suboptimal and marginal habitat in Santa Clara County. Coast redwoods have grown for decades in suboptimal (typically near riparian zones in our model) and even in marginal habitat over 300 m from streams. Many of these trees have achieved significant size despite being at lower, hotter elevations well inland of the base of the Santa Cruz Mountains. For example, the historic El Palo Alto redwood tree in Palo Alto, California, first discovered by the Portola Expedition in 1769, is located at only 20.0 m elevation and is 11 km inland from our optimal habitat’s lower limit. It is 1,083 years old, 2.3 m diameter at breast height (DBH), and 33.5 m (110 ft) tall (79). El Palo Alto is in suboptimal habitat adjacent to a seasonal creek. However, there are other large redwoods thriving in marginal habitat over 300 m distant from any nearby creeks including a heritage tree in Los Altos, California that is ∼100 years old but 2.0 m DBH and 40.5 m (133 ft) tall at 47.5 m elevation, or another heritage tree in Palo Alto that is 130 years old, 1.6 m DBH, and 38.1 m tall (125 ft) at 13.6 m elevation (79). These large trees in suboptimal and marginal habitat highlight potential additional opportunities for redwoods to sequester carbon in Santa Clara County.

However, after 100 years of growth or more some redwoods planted in suboptimal or marginal habitat may sequester less carbon as apical or crown dieback often occurs in the lower, more inland, arid conditions of the Santa Clara Valley floor. Apical dieback is highlighted by El Palo Alto which has lost one-third of its height since 1951 due to nearby creek channel incision and lowering of the water table (79).

To achieve much greater carbon offsets, the machine-learning model could be applied to identify optimal planting habitat in other California Counties. Local sampling of nearby mature redwoods would be required for ML training. In fact, nearly all of the world’s tallest forests grow in coastal, mild and stable temperature habitats, within 100 km of an ocean (25, 80). Santa Clara County is not coastal and is thus generally less favorable coast redwood habitat than counties adjacent to the Pacific coast. As expected, the model reflects that only a small portion of inland Santa Clara County, the westernmost, is optimal coast redwood habitat. Coastal California counties in coast redwoods’ historical range offer much greater opportunities to restore forest and offset carbon emissions. To illustrate the potential additional carbon sequestration opportunity, we have ordered by land area the eight California coastal counties and four inland counties in California, and the single Oregon county, in historical coast redwood range (**Table 3**).

**Table 3.**
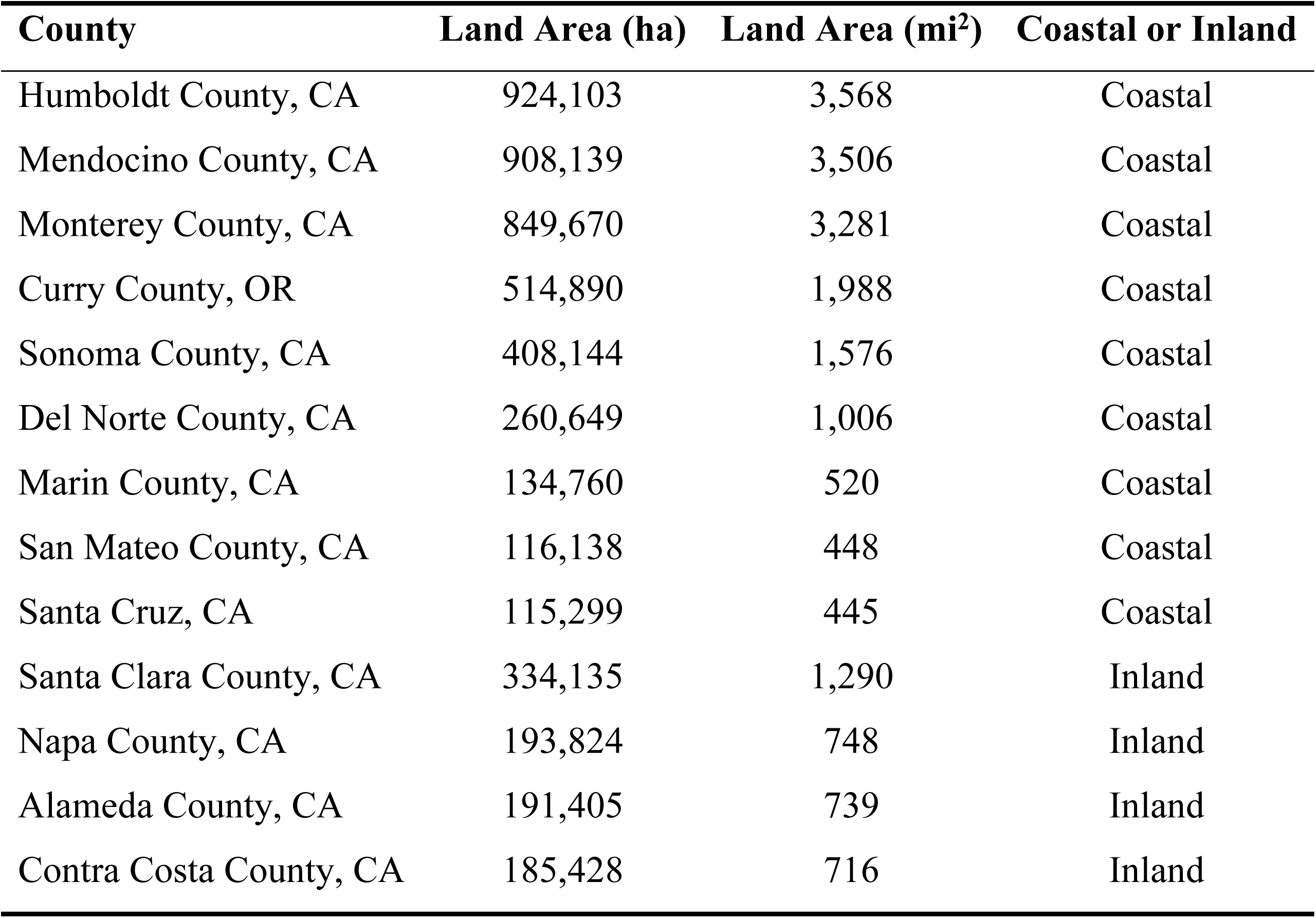
Counties in historical coast redwood range ordered by total land area. Inland counties are listed last as only small portions are likely to offer optimal redwood habitat.

Depending on the extent of deforested lands available for coast redwood reforestation, coastal California counties may possess order of magnitude higher opportunities for carbon sequestration than inland Santa Clara County as the latter only has 10.2% of its 334,135 ha (1,290mi^2^) as optimal redwood habitat. Compare this to Marin County, a relatively small coastal county but which has been modeled to have 75% of its 134,760 ha (520 mi^2^) amenable to redwoods (48). Lastly, although a warming, drier climate may be expected to contract the southern end of coast redwoods range in California (81), assisted migration of coast redwoods to coastal areas north of their historical range in Oregon [which has lost 24% of its forest cover to wildfires since 2000 (10)] may provide further opportunities to establish fire-resistant redwood forests and mitigate global warming (76,82,83). California coast redwood seedlings have been established for decades even further north in over 200 communities in Washington State, including the University of Washington Botanic Gardens Washington Park Arboretum (84).

Although it was beyond the scope of this study to address, a second opportunity to maximize carbon sequestration is to carefully manage existing coast redwood forest by focusing on two goals: reduction of fire fuel loads which act as fire stepladders that enable crown fires which may kill younger trees, and thinning of second-growth forest to promote accelerated growth rates (17, 85). Prior to the twenty-first century, high severity crown fires in coast redwood forests were rare, however, recent fire suppression and associated buildup of fuel loads has contributed to high-severity crown fires (51, 86). Prescribed and cultural burning and mechanical reductions in fire fuel loads have been shown to decrease severity of wildfires in California (87, 88). Extirpation of elk (*Cervus canadensis* ssp.), California’s native large ungulate grazer, also contributes to accumulation of fire fuel loads (89, 90), and elk have not yet been restored to their native habitat in western Santa Clara County (56, 91). Thinning of second-growth coast redwood forest, enables more rapid growth of older larger trees, as the latter acquire more wood (carbon) volume annually as a product of tree height and trunk size, and aboveground tree vigor (18). Large, older trees not only store carbon, but they also actively fix much larger amounts of carbon than smaller trees, such as plantation trees; with some very large trees sequestering as much carbon in one year as has accumulated in the entire lifetime of a mid-sized forest tree (92). Further, unlike smaller forest tree species where annual incremental additions in tree biomass decline in older age, both coast redwoods and giant sequoia acquire enlarging annual increments of wood volume until extrinsic forces cause tree death (18). Although the current study focused on identification of planting sites, careful management of existing redwood forest may produce more rapid results, with a recent global study of forest-based carbon sequestration finding 61% of the potential in protection and management of existing forests and only 39% of the potential in restoring forests lost to timber harvest (93). However, in Santa Clara County, only about 23% of the optimal redwood habitat has coniferous forest today, whereas 77% could be reforested. Therefore, both protection and careful management of existing second-growth forest as well as planting the more extensive, but unforested, optimal habitat identified here, should be pursued in parallel.

A key limitation of our model is the potential future impact of global warming, with rising temperatures and increased aridity potentially contracting the area of predicted optimal habitat. However, as discussed above, redwoods have thrived for over a century in Santa Clara County in habitat we conservatively excluded as suboptimal or even marginal. To buffer against anticipated further global warming, we have excluded suboptimal and marginal habitat from our carbon sequestration estimates. Secondly, redwoods have thrived for millennia 125 km further south in warmer, drier climates at their southern limit in southern Monterey County. Thirdly, experiments planting redwoods in drier, hotter habitats outside their historical range in California has shown success, highlighting the high genetic diversity and adaptability of the species (76, 83).

Fourthly, the elevation of our optimal habitat ranges from the summit of the Santa Cruz Mountains at 1,154 m (3,786 ft) down to 175 m (574 feet). Since temperature lapses with elevation at 6.5**°**C per 1,000 m elevation and current climate models for Santa Clara County forecast average annual temperature increases of 2.7°C [range 1.9°C to 3.2°C (3.4°F to 5.8°F)], we expect much of today’s higher elevation optimal habitat to remain satisfactory (94, 95). An advantage of the ML model is that land surface temperature is a model input variable which can be adjusted if climate predictions change. Whether the importance value of 0.16 grants sufficient weight to temperature, as well as other input variables, should be evaluated over time and in different localities. In addition, any contraction of habitat in California from higher summer temperatures may be offset by expansion of suitable habitat in Oregon and Washington from higher winter temperatures (84). The calculation of carbon sequestration may be an over- or under-estimate due to standard error; however, it is likely conservative as a recent study found that current temperature biosphere models underestimate gross primary productivity by 20% (96).

Loss of planted trees due to poor seedling survival from global warming could be a significant limitation, however our experimental seedling plots indicated high cumulative 74% survival likely due to seed selection and several practical compensatory actions. To optimize survival of new plantings, seedlings were grown from seed banks matched to the locality and elevation of the inland Santa Cruz Mountains, i.e. matched to optimal habitat. Other optimization measures included growing the one-year seedlings for an additional 19 months under close observation with initial monthly watering and to allow them to mature to age 2.5 years prior to transplantation into the wild. Once planted in the wild, they were planted in shade, planted within 25 m of an ephemeral stream, and planted at the beginning of the rainy season in November.

Further compensation for unanticipated future mortality losses could be achieved by planting seedlings at higher densities, followed by thinning if such mortality did not occur. Also, coast redwoods seedlings grown from CAL FIRE seed banks from warmer lower and more southerly locations could be planted as a form of assisted migration (83). A recent demonstration of assisted migration was successful planting oyamel (*Abies religiosa*) seedlings to higher elevations than their existing range in Mexico in anticipation of global warming (97).

A third potential limitation to our carbon sequestration estimate would be loss of planted redwoods from wildfire. Although redwoods are fire resistant, they must achieve sufficient bark thickness to protect the living cambium beneath. The trees in the 2020 CZU Fire in BBRSP were subjected to a high severity burn with 77% of canopy lost due to accumulation of high fire fuel loads (47). Despite this canopy fire, a megafire of high intensity which burned all of most trees’ leaves and needles, epicormic sprouting occurred along their trunks within months, likely fueled by carbohydrates synthesized by the trees decades earlier (24). Three years the 2020 CZU fire, a survey found that almost all coast redwoods with DBH greater than 50 cm (1.6 ft) had survived, with young sprouts all along their blackened, charred trunks (98). Nevertheless, reduction in fire fuel loads in planting areas is recommended to enable redwoods to achieve the 50 cm (1.6 ft) DBH size associated with fire resistance, estimated at 50 years in good habitat conditions (16). As California becomes increasingly fire-prone with climate change, periodic monitoring and management of planted redwood forest will be required to assure that carbon stock additionality is achieved (6, 15).

Reforestation of over 26,051 ha (101 square miles) with redwood forest will require access to land, seedling supply, and labor. Fortunately, 31% of the County’s lands are already conserved and 78% of optimal habitat falls on these lands (Fig 5). Local governmental organization landholders for most of this land include Santa Clara County Parks, Santa Clara County Open Space Authority, Santa Clara Valley Habitat Agency, and Midpeninsula Open Space Preserves District, and large private landholders include Peninsula Open Space Trust, Save the Redwoods, and Sempervirens Fund. All have been contacted by the coauthors and are supportive of the concept of restoration and management of coast redwood forest to sequester atmospheric carbon. Private and public entities, such as the North Santa Clara Resource Conservation District, may obtain low-cost coast redwood seedlings grown from the California seed bank by CAL FIRE L. A. Moran Reforestation Center (64). Santa Clara County-based tree planting organizations include the public and private entities such as the California Conservation Corps, Grassroots Ecology, and Our City Forest. In sum, the resources and organizational components required for a county-level redwood planting initiative are in place in Santa Clara County, but will require coordination and commitment, accompanied by careful planning and prioritization to achieve significant and enduring carbon offsets.

## CONCLUSIONS

Along with reduction in burning fossil fuels and careful management and preservation of existing forests, planting new coast redwood forests provides a significant opportunity to offset greenhouse gas emissions (93). Arguments against tree planting to sequester carbon center on two points: first that individual trees have insufficient biomass to significantly mitigate greenhouse gas emissions and second that the trees do not live long enough to maintain significant sequestered carbon (99). Those assertions are upended when tree species such as coast redwoods are planted, due to their rapid and enduring growth rate, ultra-high biomass, exceptional longevity, and fire resistance. Coupling proper tree species selection with identification of optimal habitat via a machine-learning model, we find that tree planting of coast redwoods in a portion of a single California county could provide significant statewide carbon offsets.

## ACKNOWLEDGMENTS

First, the authors thank Todd Steiner, Preston Brown, and Elizabeth Villano of the Turtle Island Restoration Network team, whose original idea to model GIS maps of coast redwood planting habitat inspired us to apply a machine learning model to build upon and scale their approach. Second, we thank the North Santa Clara Resource Conservation District for research grant funding. We also appreciate Sonia Morris and Joanna Nelson, of the Save the Redwoods League, and Professor Robert Van Pelt, University of Washington, who provided their respective maps of the historical range of coast redwoods. Lastly, we thank Kuldeep Singh and Katherine Bolte of the CAL FIRE Lewis. A. Moran Reforestation Center for granting 250 one-year old coast redwood seedlings for our experimental plantings.

